# A high throughput screening assay for inhibitors of SARS-CoV-2 pseudotyped particle entry

**DOI:** 10.1101/2021.10.04.463106

**Authors:** Miao Xu, Manisha Pradhan, Kirill Gorshkov, Jennifer D. Petersen, Min Shen, Hui Guo, Wei Zhu, Carleen Klumpp-Thomas, Sam Michael, Misha Itkin, Zina Itkin, Marco R. Straus, Joshua Zimmerberg, Wei Zheng, Gary R. Whittaker, Catherine Z. Chen

## Abstract

Effective small molecule therapies to combat the SARS-CoV-2 infection are still lacking as the COVID-19 pandemic continues globally. High throughput screening assays are needed for lead discovery and optimization of small molecule SARS-CoV-2 inhibitors. In this work, we have applied viral pseudotyping to establish a cell-based SARS-CoV-2 entry assay. Here, the pseudotyped particles (PP) contain SARS-CoV-2 spike in a membrane enveloping both the murine leukemia virus (MLV) gag-pol polyprotein and luciferase reporter RNA. Upon addition of PP to HEK293-ACE2 cells, the SARS-CoV-2 spike protein binds to the ACE2 receptor on the cell surface, resulting in priming by host proteases to trigger endocytosis of these particles, and membrane fusion between the particle envelope and the cell membrane. The internalized luciferase reporter gene is then expressed in cells, resulting in a luminescent readout as a surrogate for spike-mediated entry into cells. This SARS-CoV-2 PP entry assay can be executed in a biosafety level 2 containment lab for high throughput screening. From a collection of 5,158 approved drugs and drug candidates, our screening efforts identified 7 active compounds that inhibited the SARS-CoV-2-S PP entry. Of these seven, six compounds were active against live replicating SARS-CoV-2 virus in a cytopathic effect assay. Our results demonstrated the utility of this assay in the discovery and development of SARS-CoV-2 entry inhibitors as well as the mechanistic study of anti-SARS-CoV-2 compounds. Additionally, particles pseudotyped with spike proteins from SARS-CoV-2 B.1.1.7 and B.1.351 variants were prepared and used to evaluate the therapeutic effects of viral entry inhibitors.

## Introduction

The current global pandemic of coronavirus disease 2019 (COVID-19) has escalated rapidly over the last year into a disaster affecting all aspects of human life. While effective vaccines are now available, specific antiviral treatment for COVID-19 is still an unmet and urgent need ^1^. Because the severe acute respiratory syndrome coronavirus 2 (SARS-CoV-2) that causes COVID-19 is a biosafety level-3 (BSL-3) virus, specific and high containment laboratory conditions are required to handle live virus experiments. However, most laboratories conducting high throughput screening (HTS) of compounds are limited to BSL-2 assays, a significantly limitation to lead discovery efforts.

Although virtual screens using in silico modeling technologies have been employed by many laboratories to search for anti-SARS-CoV-2 compounds, the anti-SARS-CoV-2 activities of these compounds must be evaluated in physical experiments. For example, a recent drug repurposing screen using a SARS-CoV-2 cytopathic effect (CPE) assay has confirmed the anti-SARS-CoV-2 activities of several compounds reported by virtual screening studies ^2^. The lack of SARS-CoV-2 assays suitable for BSL-2 laboratory settings that are accessible to most researchers may contribute to missing experimental confirmation of virtual screening hits. Therefore, more BSL-2 compatible SARS-CoV-2 compound screening assays are needed for COVID-19 drug development.

The spike (S) proteins of coronaviruses are responsible for host receptor binding and fusion of viral bilayer membranes with host cell membranes to release the viral genome in the cytoplasm ^3^. To alleviate SARS-CoV-2 biosafety issues, several viral pseudotyping systems have been reported for the SARS-CoV-2-S protein, including the murine leukemia virus (MLV) pseudotyping method ^4, 5^. We have recently employed MLV pseudotyped particle (PP) viral entry assays for SARS-CoV-S and MERS-CoV-S to screen compound libraries for viral entry inhibitors in a BSL-2 setting ^4^. These viral entry assays utilize PPs that consist of bilayer membranes containing spike proteins and package luciferase reporter RNA instead of a viral genome. The binding of PPs to receptors on the host cell plasma membrane, such as ACE2 for SARS-CoV and SARS-CoV-2, and dipeptidyl peptidase-4 (DPP4) for MERS-CoV, mediates entry of PPs into host cells. Upon entry, the luciferase reporter RNA is expressed, and after the introduction of the luciferase substrate, the luciferase enzymatic reaction produces a luminescence signal can be detected and used as an indicator of viral entry into cells.

Here we report a BSL-2 compatible SARS-CoV-2 PP assay that has been characterized and validated in a drug repurposing screen of over 5,000 compounds. Our results demonstrate that this SARS-CoV-2-S PP entry assay is robust for HTS and can be used for lead discovery and lead optimization. In addition, this assay can be used to determine whether the mechanism of action of anti-SARS-CoV-2 compounds found in other phenotypic screening assays is due to the inhibition of viral entry. We have also used the PP entry assay to evaluate the therapeutic effects of entry inhibitors towards the SARS-CoV-2 spike variants B.1.1.7 and B.1.351. Thus spike proteins from newly emerging SARS-CoV-2 variants can be rapidly evaluated with this surrogate assay for viral entry.

## Materials and Methods

### Materials

Dulbecco’s Modified Eagle’s Medium (DMEM) (11965092), OPTI-MEM reduced serum medium (11058021), Pen/Strep (15140), TrypLE (12604013), PBS -/-(w/o Ca2+ or Mg2+) (10010049), Lipofectamine 3000 Transfection Reagent (L3000015), HCS Cell Mask Green (H32714), goat anti-mouse AlexaFluor 647 (A28175), and Hoechst 33342 (H3570) were purchased from ThermoFisher. Hyclone FBS (SH30071.03) was obtained from GE Healthcare. Microplates including white tissue culture treated 96-well plates (655090), black μClear 96-well plates (655083), white tissue culture-treated 384-well plates (781073), and white tissue-culture treated 1536-well plates (789173-F) were purchased from Greiner Bio-One. BrightGlo Luciferase Assay System (E2620) was obtained from Promega. ATPLite Luminescence Assay kit was purchased from PerkinElmer (6016949). Cell Staining Buffer (420201) was purchased from BioLegend. Paraformaldehyde (PFA) was purchased from Electron Microscopy Sciences (15714-S). Mouse-anti-firefly luciferase antibody was purchased from Santa Cruz (sc-74548). SARS-CoV-2 spike antibody was purchased from Sino Biological (40589-T62).

### Pseudotyped particle (PP) generation

SARS-CoV-2-S pseudotyped particles (PPs) as well as the control PPs (VSV-G PP and bald PP were custom produced by the contract research organization (CRO) Codex Biosolutions (Gaithersburg, MD). PP production was based on previously reported methods using a murine leukemia virus (MLV) pseudotyping system ^6, 7^. The SARS-CoV2-S construct with Wuhan-Hu-1 sequence (BEI #NR-52420) was C-terminally truncated by 19 amino acids to increase incorporation into PPs ^5^. For pseudotyping SARS-CoV-2-S variants, B1.1.7 (alpha) variant bearing mutations del69-70, del144, N501Y, A570D, D614G, P681H, T716I, S982A, and D1118H, and B.1.351 (beta) variant bearing mutations L18F, D80A, D215G, del242-244, K417N, E484K, N501Y, D614G, and A701V, were combined with C-terminal 19 amino acid truncation for pseudotyping.

The SARS-CoV-2 PPs were produced as followings. 7E6 HEK293T cells were seeded in a 10 cm^2^ dish in 16 ml culture medium (DMEM, 10% FBS) without antibiotics that were incubated for 24 hr at 37 °C with 5% CO_2_. Lipofectamine 3000 Transfection Reagent (ThermoFisher) was used for transfection. The DNA solution was prepared by diluting 8 μg of pTG-Luc, 6 μg of pCMV-MLVgag-pol, and 6 μg of pcDNA-SARS-CoV-2-S(ΔC19) in 1 ml of Opti-MEM medium (ThermoFisher). Then, 40 μl of P3000 reagent was added to the DNA solution. The DNA-P3000 mixture was further mixed with 40 μl of Lipofectamine 3000 reagent pre-diluted in 1 ml Opti-MEM medium. After a 10 min incubation at room temperature, the mixture of DNA/P3000/Lipofectamine complex was added to cells in 10-cm dish, and incubated at 37°C with 5% CO_2_ for 48 hr. The supernatant was collected and centrifuged at 290 x g for 7 min to remove cell debris. The supernatant was then passed through a 0.45 μm PVDF Syringe Filter (Santa Cruz, Cat# sc-358814). The final supernatants collected from 10 cm^2^ dishes were pooled, aliquoted, and stored at 4°C for short term use (up to 14 days), or at -80°C for long term storage.

### Pseudotyped particle (PP) entry assay in 1536-well format

Expi293F cell line with stable expression of human ACE2 (HEK293-ACE2) cells ^8^ were seeded in white, solid bottom 1536-well microplates (Greiner BioOne) at 1500 cells/well in 2 μL/well medium (DMEM, 10% FBS, 1x L-glutamine, 1x Pen/Strep, 1 ug/ml puromycin), and were incubated at 37 °C with 5% CO2 overnight (∼16 hr). Compounds were titrated at a 1:3 dilution in DMSO and dispensed via pintool (Waco Automation, San Diego, CA) at 23 nL/well to assay plates. Cells were incubated with the test compounds for 1 h at 37 °C with 5% CO2, before 2 μL/well of PPs were added. The plates were then spinoculated by centrifugation at 1500 rpm (453 x g) for 45 min at room temperature, and incubated for 48 hr at 37 °C with 5% CO2 to allow cell entry of PPs and expression of luciferase reporter. After the incubation, the supernatant was removed with gentle centrifugation using a Blue Washer (BlueCat Bio). Then 4 μL/well of Bright-Glo Luciferase detection reagent (Promega) was added to assay plates and incubated for 5 min at room temperature. The luminescence signal was measured using a PHERAStar plate reader (BMG Labtech). Data was normalized with wells inoculated with SARS-CoV-2-S PPs as 100%, and wells inoculated with bald PPs (no spike protein) as 0%.

### Pseudotyped particle (PP) entry assay in 384-well format

HEK293-ACE2 cells ^8^ were seeded in white, solid bottom 384-well microplates (Greiner BioOne) at 6000 cells/well in 10 μL/well medium, and were incubated at 37 °C with 5% CO_2_ overnight (∼16 hr). After an overnight incubation, 5 μl/well of compound solution or medium was added and incubated at 37°C for 1 hr, followed by the addition of 15 μL/well of PPs. The plates were then spinoculated by centrifugation at 1500 rpm (453 xg) for 45 min, and incubated for 48 hr at 37 °C with 5% CO_2_ to allow cell entry of PPs and expression of luciferase reporter. After the incubation, the supernatant was removed with gentle centrifugation using a Blue Washer (BlueCat Bio). Then 20 μL/well of Bright-Glo Luciferase detection reagent (Promega) was added to assay plates and incubated for 5 min at room temperature. The luminescence signal was measured using a PHERAStar plate reader (BMG Labtech). Data was normalized with wells inoculated with fusion glycoprotein bearing PPs as 100%, and wells inoculated with bald PPs (no spike protein) as 0%.

### ATP content cytotoxicity assay in 1536-well format

HEK293-ACE2 cells were seeded in white, solid bottom 1536-well microplates (Greiner BioOne) at 2000 cells/well in 2 μL/well medium, and incubated at 37 °C with 5% CO2 overnight (∼16 hr). Compounds were titrated 1:3 in DMSO and dispensed via pintool at 23 nL/well to assay plates. Cells were incubated with test articles for 1 h at 37 °C with 5% CO2, before 2 μL/well of media was added. The plates were then incubated at 37 °C for 48 h at 37C with 5% CO2. After incubation, 4 μL/well of ATPLite (PerkinElmer) was added to assay plates and incubated for 15 min at room temperature. The luminescence signal was measured using a Viewlux plate reader (PerkinElmer). Data was normalized with wells containing cells as 100%, and wells containing media only as 0%.

### Luciferase immunofluorescence and high-content imaging

Cells were seeded at 15,000 cells in 100 μL/well media in 96-well assay plates, and incubated at 37 °C, 5% CO_2_ overnight (∼16 hr). Supernatant was removed, and 50 μL/well of PP was added. Plates were spinoculated at 1500 rpm (453 xg) for 45 min, incubated for 2 hr at 37 °C with 5% CO_2_, then 50 μL/well of growth media was added. The plates were incubated for 48 hr at 37 °C with 5% CO_2_. Media was aspirated, and cells were washed once with 1X PBS (ThermoFisher). Cells were then fixed in 4% PFA (EMS) in PBS containing 0.1% BSA (ThermoFisher) for 30 min. at room temperature. Plates were washed three times with 1X PBS, then blocked and permeabilized with 0.1% Triton-X 100 (ThermoFisher) in Cell Staining Buffer (Biolegend) for 30 min. Permeabilization/blocking solution was removed, 1:1000 primary mouse-anti-luciferase antibody (Santa Cruz) was added, and incubated overnight at 4 °C. Primary antibody was aspirated and cells were washed three times with 1X PBS. 1:1000 secondary antibody goat-anti-mouse-AlexaFluor 647 (ThermoFisher) was added for 1 hr in Cell Staining Buffer. Cells were washed three times, and stained with 1:5000 Hoechst 33342 (ThermoFisher) and 1:10000 HCS Cell Mask Green (ThermoFisher) for 30 min. before three final 1X PBS washes. Plates were sealed and stored at 4 °C prior to imaging.

Plates were imaged on the IN Cell 2500 HS automated high-content imaging system. A 20x air objective was used to capture nine fields per well in each 96 well plate. Cells were imaged with the DAPI, Green, and FarRed channels. Images were uploaded to the Columbus Analyzer software for automated high-content analysis. Cells were first identified using the Hoechst 33342 nuclear stain in the DAPI channel. Cell bodies were identified using the HCS Cell Mask stain in the green channel using the initial population of nuclei regions of interests. Intensity of the FarRed channel indicating luciferase expression was measured, and a threshold was applied based on the background of the negative control. Average values, standard deviations, and data counts were generated using pivot tables in Microsoft Excel and data was plotted in Graphpad Prism.

### Negative stain transmission electron microscopy and immunogold labeling

All reagents were obtained from Electron Microscopy Sciences, unless otherwise specified. SARS2-CoV-2 PPs were adhered to freshly glow discharged, formvar and carbon coated, 300-mesh gold grids by inverting grids on 5 μl drops of supernatant containing PPs for 1 min, and then rinsed by transferring the grids across 3 droplets of filtered PBS. Grids were covered during the incubation steps to prevent evaporation. Next, grids were incubated on drops of blocking solution containing 2% BSA (Sigma) in PBS for 10 min. Primary antibody to SARS-CoV-2 spike (Sino Biological, 40589-T62), was diluted 1:20 in blocking solution and grids were transferred to drops of primary antibody for 30 minutes. Then, grids were transferred across two drops of blocking solution for 10 minutes before incubation of with 10 nm gold-conjugated goat-α-rabbit secondary antibody diluted 1:20 in blocking solution for 30 min. Finally, grids were rinsed with 3 drops of PBS and 3 drops of filtered distilled water before being placed on a drop of 1% aqueous uranyl acetate negative staining solution for 30 seconds, after which grids were blotted with filter paper, allowing a thin layer of uranyl acetate to dry on the grid. Grids were observed using a ThermoFisher Tecnai T20 transmission electron microscope operated at 200 kV, and images were acquired using an AMT NanoSprint1200 CMOS detector (Advanced Microscopy Techniques).

### Drug repurposing screen and data analysis

The NCATS pharmaceutical collection (NPC) was assembled internally, and contains 2,678 compounds, which include drugs approved by US FDA and foreign health agencies in European Union, United Kingdom, Japan, Canada, and Australia, as well as some clinical trialed experimental drugs ^9^. The NCATS Mechanism Interrogation Plate (MIPE) 5.0 library contains 2,480 mechanism based bioactive compounds, targeting more than 860 distinct mechanisms of action. The compounds were dissolved as 10 mM DMSO stock solutions, and titrated at 1:5 for primary screens with 4 concentration points. The SARS-CoV-2-S PP entry assay and ATP content cytotoxicity assay in HEK293-ACE2 cells, were used to screen the NPC and MIPE libraries in parallel. The primary screens assayed four compound concentrations at the single point. Hit compounds were cherry picked, titrated at 1:3 for follow up assays with 11 concentrations, in duplicates. Hit compounds were confirmed in SARS-CoV-2-S PP entry assay and cytotoxicity assay, as well as counter screened in a VSV-G PP entry assay in the same cell line.

A customized software developed in house at NCATS ^10^ was used for analyzing the primary screen data. Half-maximal efficacious concentration (EC_50_) and half-maximal cytotoxicity concentration (CC_50_) of compounds were calculated using Prism software (GraphPad Software, San Diego, CA).

### SARS-CoV-2 cytopathic effect (CPE) assay

SARS-CoV-2 CPE assay was conducted at a contract research organization Southern Research Institute (Birmingham, AL) ^2^. Briefly, compounds were titrated in DMSO and acoustically dispensed into 384-well assay plates at 60 nL/well. Cell culture media (MEM, 1% Pen/Strep/GlutaMax, 1% HEPES, 2% HI FBS) was dispensed at 5 μL/well into assay plates and incubated at room temperature. Vero E6 (selected for high ACE2 expression) was inoculated with SARS-CoV-2 (USA_WA1/2020) at 0.002 M.O.I. in media and quickly dispensed into assay plates as 4000 cells in 25 μL/well. Assay plates were incubated for 72 hr at 37 °C, 5% CO_2_, 90% humidity. Then, 30 μL/well of CellTiter-Glo (Promega) was dispensed, incubated for 10 min at room temperature, and luminescence signal was read on an EnVision plate reader (PerkinElmer). An ATP content cytotoxicity assay was conducted with the same protocol as CPE assay, without the addition of SARS-CoV-2 virus.

## Results

### SARS-CoV-2-S pseudotyped particle entry assay design

Production of PPs was done largely following previously reported protocols for SARS and MERS PPs with minor changes ^6, 7^. Three plasmids were used to co-transfect HEK293 cells to produce the pseudotyped particles (PPs) containing capsid protein of murine leukemia virus (MLV), SARS-CoV-2 spike protein, and luciferase RNA (**Figure 1a**). Two of the three plasmids, pCMV-MLVgag-pol and and pTG-Luc were same as previously described ^6^, and the third plasmid for the SARS-CoV-2 spike protein was newly generated. The PPs produced by the co-transfection carry SARS-CoV-2 spike protein on their bilayer surface, and package firefly luciferase RNA instead of the SARS-CoV-2 genome. Thus, the PPs are only capable of a single round of entry into host cells via ACE2 receptor-mediated entry, and cannot replicate in cells. PP entry is complete when membrane fusion and luciferase RNA is released into the cell. After a 48 hr incubation period to allow for luciferase expression, the amount of PP entry could be read out via a luciferase enzyme assay with a luminescent signal (**Figure 1b**).

**Figure 1.**
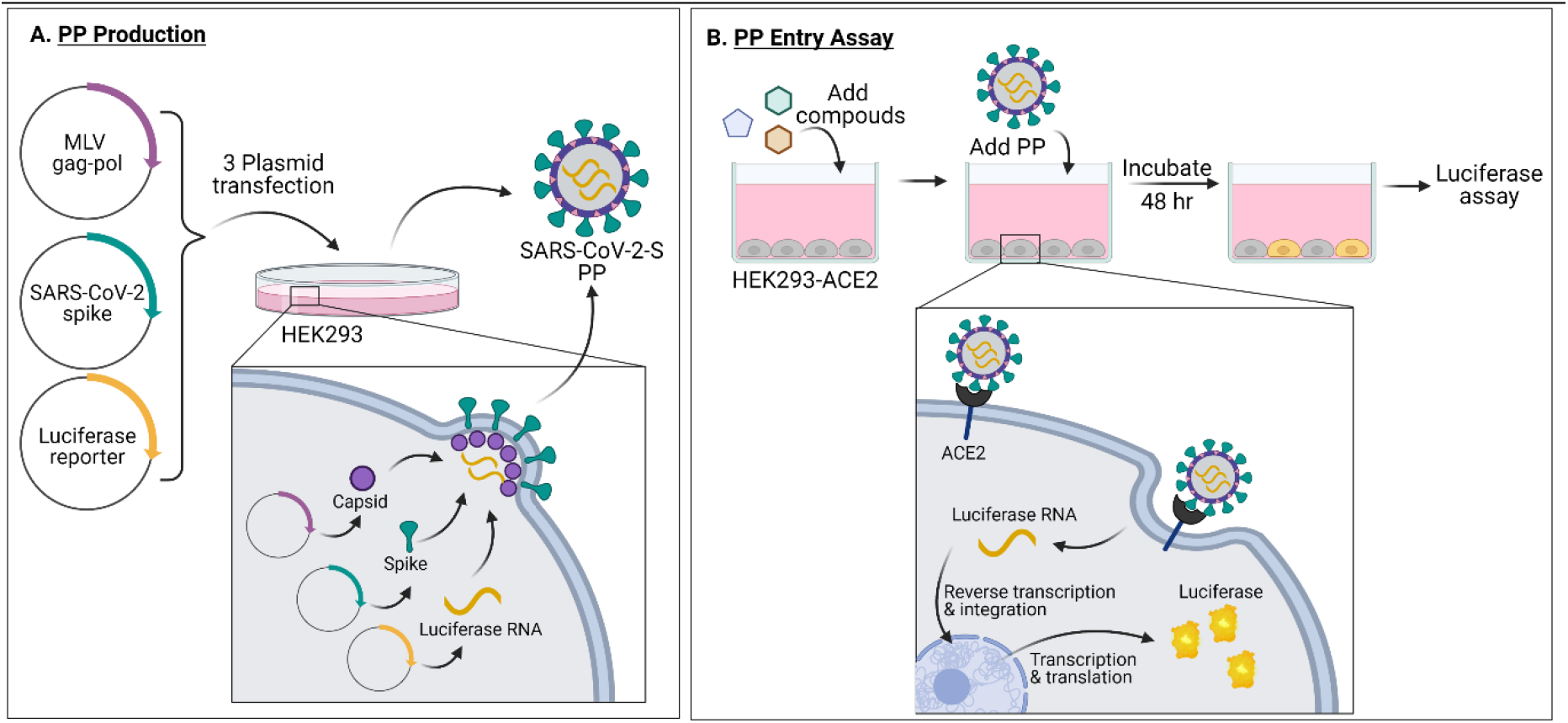
(a) PP production and (b) PP entry assay schematics. Made with Biorender.com.

### SARS-CoV-2 PP entry assay optimization and miniaturization

Binding of SARS-CoV-2 spike protein to the ACE2 receptor on cell membrane is the first step of viral entry into cells. An Expi293F cell line stably expressing human ACE2 (HEK293-ACE2) was used for this SARS-CoV-2 PP assay ^8^. After inoculation, the PPs entered cells and released luciferase RNA that resulted in expression of luciferase reporter enzyme. The luminescence signal, an indicator of the entry of SARS-CoV-2 into cells, was then detected. Cells treated with bald PPs, i.e. PPs lacking fusion glycoproteins, were used as a basal signal control. PPs pseudotyped with vesicular stomatitis virus glycoprotein G (VSV-G) were used as a control in PP production and entry assays. VSV-G is a class III fusion protein that binds host LDL receptor family members, and mediates cell entry of PPs pseudotyped with VSV-G. It is commonly used as a control in viral pseudotyping due to its wide range of cellular tropism ^11^. The entry assay with VSV-G PPs was used to eliminate entry inhibitors in our compound screens operating via the MLV components of the PP, rather than the SARS-CoV-2 components. As shown in **Figure 2a**, HEK293-ACE2 cells inoculated with VSV-G PPs showed a signal-to-basal ratio (S/B) of 776-fold compared with the basal signal generated by bald PPs that do not contain fusion glycoproteins. Bald PPs produced a slight increase in luciferase signal when compared with no PP medium control, with RLU values of 4.7 and 2.7 for bald PP and no PP, respectively (**Figure 2a**). Therefore, inoculation with bald PP was used as a negative control in all experiments. Consistent with previous reports, we found that cells treated with PPs bearing SARS-CoV-2-S with C-terminal 19 amino acid truncation (ΔC19) showed enhanced S/B ratio than full length (FL) spike ^5^, with S/B ratio of 1178 and 47.8 fold for PPs with ΔC19 spike and FL spike, respectively (**Figure 2a**). Therefore, all subsequent PP experiments were carried out with the ΔC19 spike construct, which henceforth will be referred to as the SARS-CoV-2-S PP.

**Figure 2.**
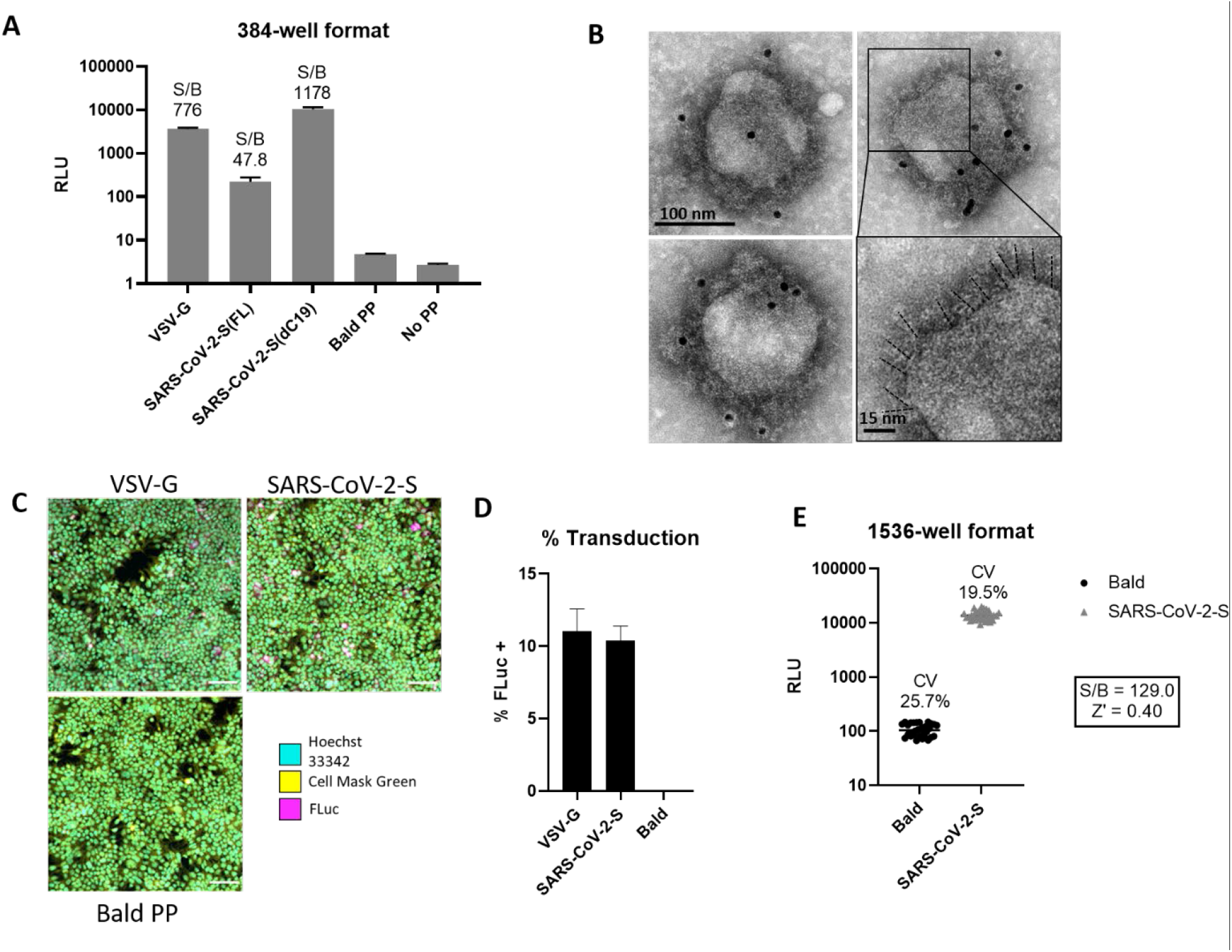
Characterization and optimization of SARS-CoV-2-S PP entry assay. (a) PP entry test in HEK293-ACE2 cells in 384-well format. Average relative luminescence unit (RLU) for bald PP (no glycoprotein) was used for signal-to-basal (S/B) calculations. (b) EM of SARS-CoV-2-S(ΔC19) PP with immunogold labeling for SARS-CoV-2 spike. Bottom right shows enlarged area with putative spikes highlighted (dashed lines). (c) Immunofluorescence staining of luciferase in PP transduced HEK293-ACE2 cells. (d) Quantitation of luciferase positive cells by immunofluorescence staining. (e) Miniaturization of PP entry assay in 1536-well format.

To confirm the expression and incorporation of SARS-CoV-2 spike proteins in the PPs, we carried out negative stain transmission electron microscopy with immunogold labeling of spike. The electron micrographs showed the presence of a spiky fringe decorating the surface of PPs that labeled with immunogold directed against the SARS-CoV-2 spike protein (**Figure 2b**). To understand the level of PP transduction, we conducted immunofluorescent staining of the luciferase enzyme in the HEK293-ACE2 cells treated with the PPs and found a significantly greater expression of luciferase reporter proteins in the SARS-CoV-2-S and VSV-G PPs treated cells compared to bald PP treatment (**Figure 2c**). These results correlate well with whole-well based luciferase assay signals (**Figure 2a**). Together, these results indicate a significant increase in luciferase luminescence after inoculating HEK293-ACE2 cells with PPs carrying SARS-CoV-2-S ΔC19.

We then miniaturized the SARS-CoV-2-S PP entry assay to the 1536-well plate format for HTS by proportionally reducing the quantity of cells and reagents (**Table 1**). A seeding density of 1500 cells per well resulted in approximately 80% confluence after an overnight incubation, which is the standard confluence for pseudovirus assays. The S/B ratio was 129-fold, CV was 19.5%, and Z’ factor was 0.40 in the 1536-well plate (**Figure 2e**). These values indicated that this assay was acceptable for HTS.

**Table 1.**
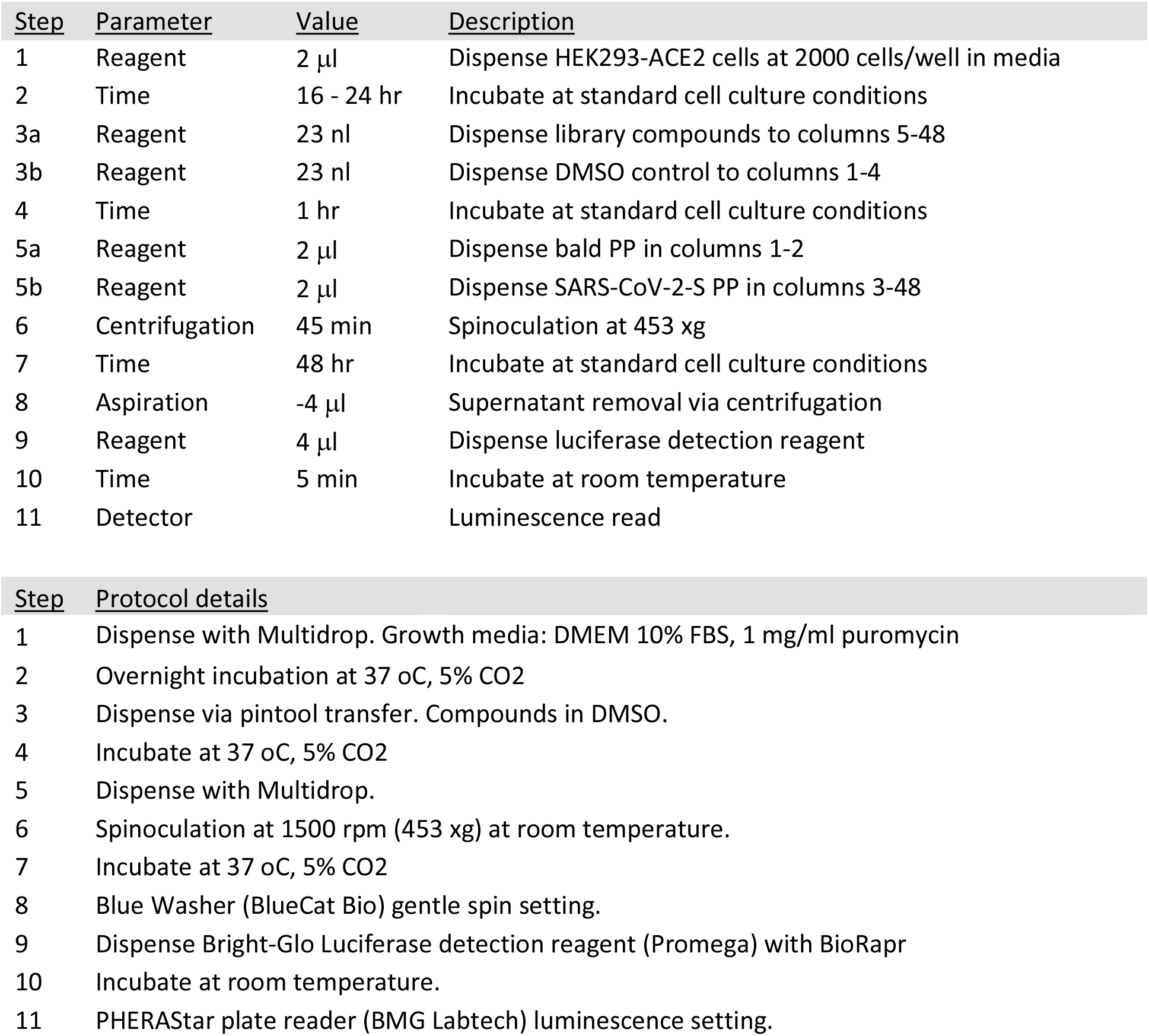
qHTS protocol

### Quantitative high throughput screening (qHTS) using the SARS-CoV-2 PP assay

A qHTS was performed using the optimized SARS-CoV-2-S PP entry assay in the 1536-well plate format. A total of 5,158 compounds, containing approved drugs, drug candidates, and mechanism-based bioactive compounds were screened at four assay concentrations (0.46, 2.30, 11.5, 57.5 μM). Due to the fact that this was a signal decrease assay and required a 48 hr incubation, cytotoxic compounds could have appeared as false positives by reducing luciferase luminescence because of cell death. Therefore, an ATP content cell viability assay was carried out in parallel with the same cells and assay protocol, without the addition of PP, to filter out the cytotoxic compounds (**Figure 3**).

**Figure 3.**
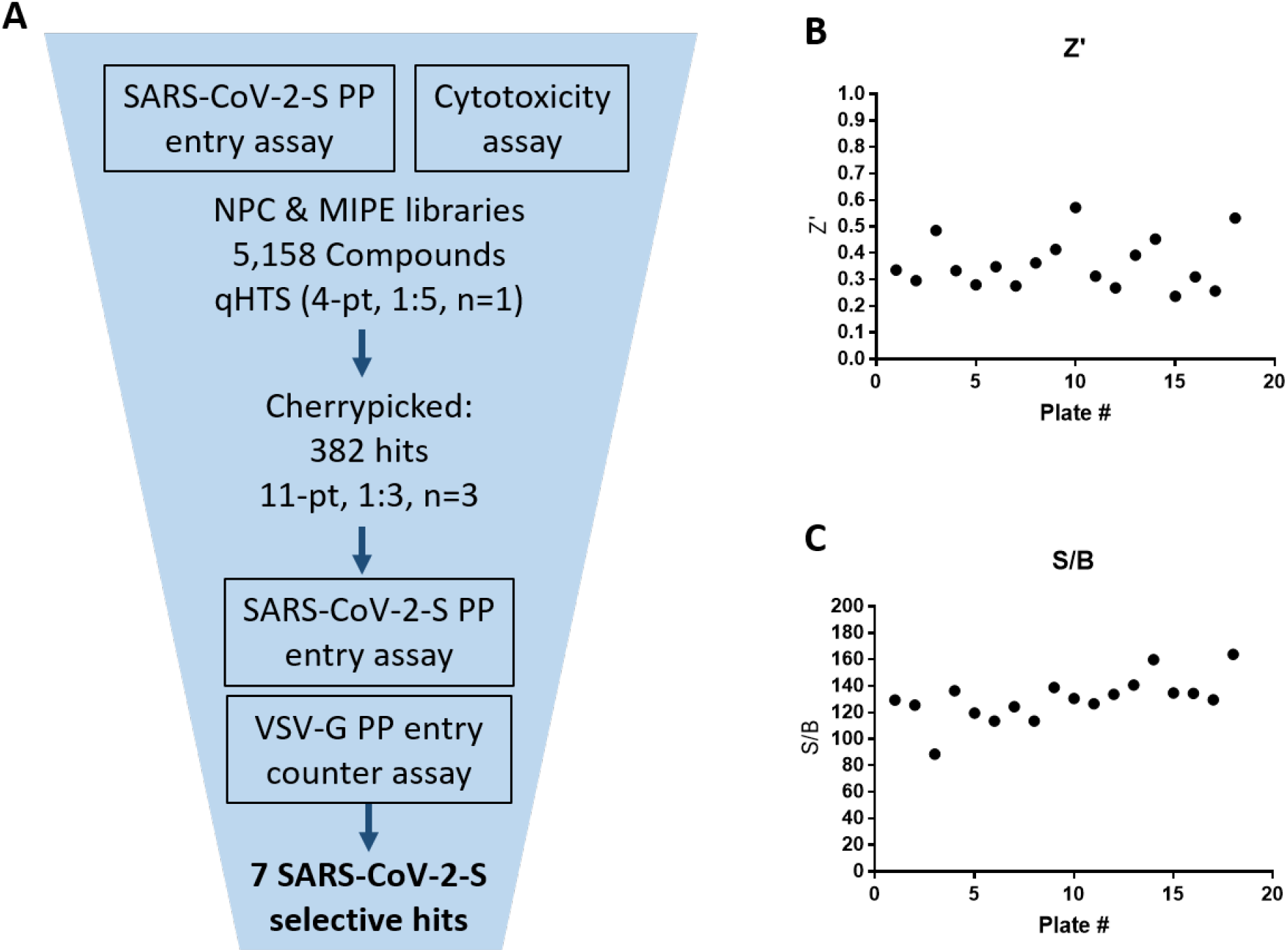
qHTS summary. (a) qHTS and follow up schematic. (b) qHTS plate z’ factors. (c) qHTS plate S/B.

The primary screening data was analyzed using an in-house curve fitting software and the primary hits were selected based on a cut-off values of EC_50_ < 10 μM and maximal inhibition > 65% with a delta AUC (area-under-the-curve difference between PP assay and cytotoxicity counter-screen) greater than 60. A set of 382 compounds were selected as the primary hits that were freshly QC’ed and retested in the SARS-CoV-2-S PP entry assay using an 11-point concentration titration (1:3 dilution). In addition, a VSV-G PP entry assay was employed as an additional counter-screen to eliminate false positive hits that inhibit luciferase expression or luciferase enzymatic activity instead of spike-mediated cell entry. From these three confirmation and counter-screen assays, seven specific inhibitors of SARS-CoV-2 entry were identified that show >10-fold selectivity between SARS-CoV-2-S and VSV-G PP entry assays (**Figure 4**). All data from the primary screen and follow up assays were publicly posted on PubChem (**Table 2**), as well as the NCATS OpenData Portal (https://opendata.ncats.nih.gov/covid19/).

**Table 2.** Public datasets. Primary screen and follow up assay data can be found at https://pubchem.ncbi.nlm.nih.gov/ under the following assay IDs (AIDs).

| AID | # of Compounds | Concentration response format | Assay |
| --- | --- | --- | --- |
| 1645846 | 5,158 | 4-pt, 1:5 | SARS-CoV-2-S PP entry |
| 1645845 | 5,158 | 4-pt, 1:5 | Cytotoxicity |
| 1645844 | 382 | 11-pt, 1:3 | SARS-CoV-2-S PP entry |
| 1645847 | 382 | 11-pt, 1:3 | VSV-G PP entry |

**Figure 4.**
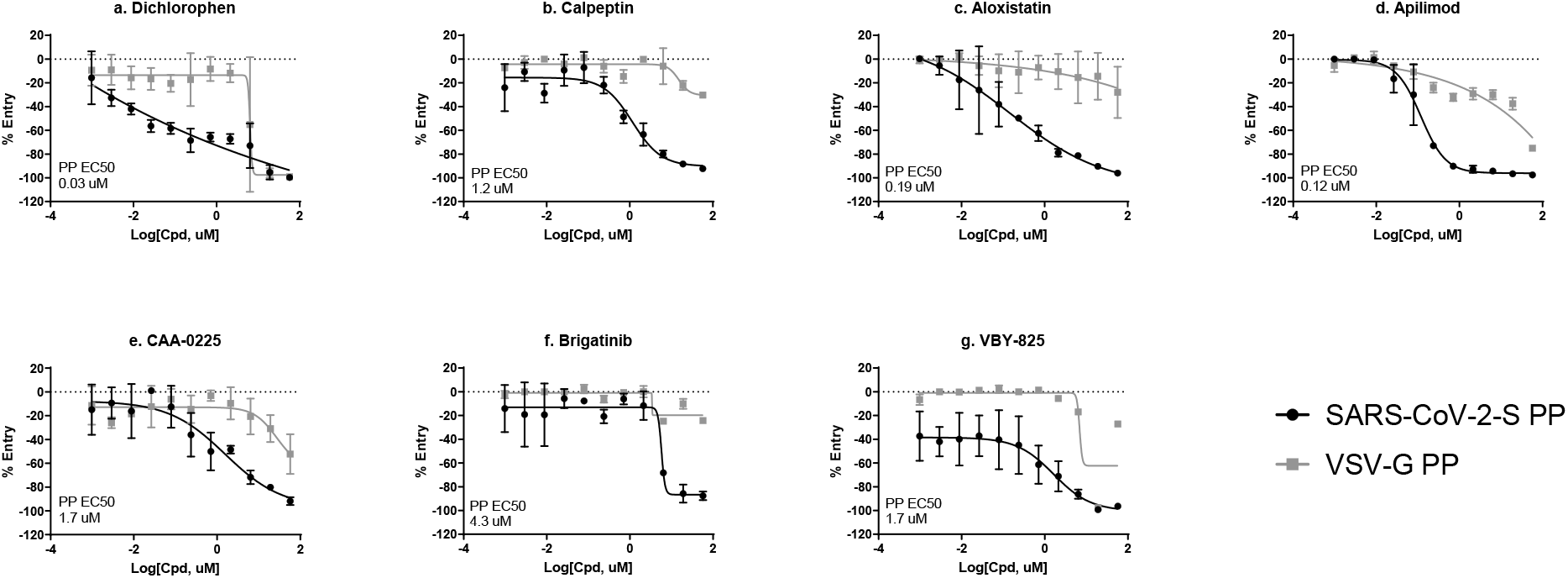
Top 7 SARS-CoV-2-S PP selective inhibitors in HEK293-ACE2 cells. (a) dichlorophen, (b) calpeptin, (c) aloxistatin, (d) apilimod, (e) CAA-0225, (f) brigatinib, and (g) VBY-825.

### Hit confirmation in a SARS-CoV-2 cytopathic effect (CPE) assay

To further confirm the entry inhibition activity, the seven selective entry inhibitors were tested in a SARS-CoV-2 live virus infection assay, by measuring cytopathic effect (CPE) of SARS-CoV-2 in VeroE6 cells ^2^. The results in this live virus assay confirmed that six compounds showed a protective effect against the cell death caused by the virus (**Figure 5**), while the activity dichlorophen was not confirmed in the CPE assay, possibly due to cell line differences.

**Figure 5.**
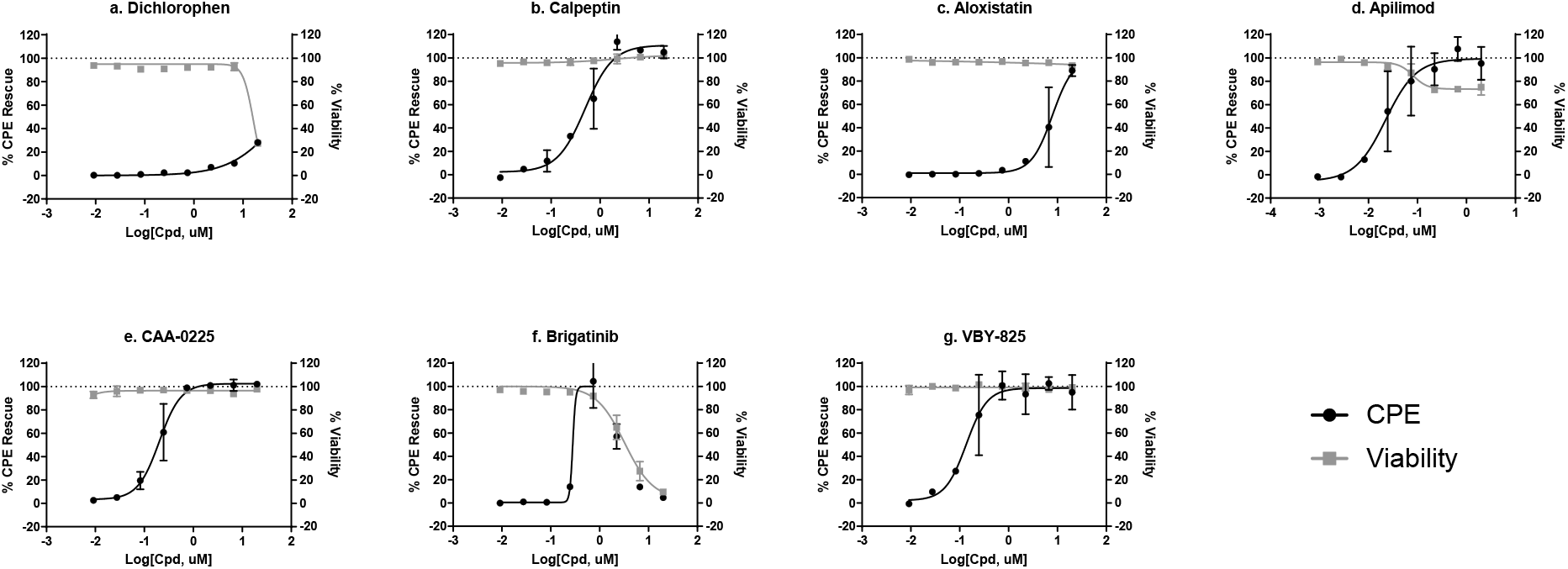
CPE rescue activity of top 7 SARS-CoV-2-S PP selective inhibitors (a) dichlorophen, (b) calpeptin, (c) aloxistatin, (d) apilimod, (e) CAA-0225, (f) brigatinib, and (g) VBY-825.

In conclusion, six compounds were identified as the specific inhibitors against viral entry into host cells mediated by the SARS-CoV-2 spike protein. Among these compounds, the SARS-CoV-2 CPE rescue was confirmed for six compounds.

### SARS-CoV-2 PP assay with mutations from the B.1.1.7 and B.1.351 variants

To evaluate the applicability of this PP entry assay to spike variants, variant spike proteins from the B.1.1.7 (alpha) and B.1.351 (beta) strains were used to pseudotype MLV PPs. The 6 entry inhibitors that were confirmed in the CPE assay were tested in 3 SARS-CoV-2-S PP entry assays, with the original Wuhan-Hu-1 spike sequence, and two variants of concern B.1.1.7 and B.1.351 spike sequences, as well as an ATP content cytotoxicity assay (**Figure 6**). The potency and efficacy of each of the six compounds in the PP entry assays were not significantly affected by these spike variants. All but one compound had EC50’s that were within 3-fold for each variant. Apilimod showed EC_50_’s of 9.2 nM, 1.3 nM and 4.7 nM in Wuhan-Hu-1, B.1.1.7 and B.1.351 SARS-CoV-2-S PP entry assays, with a 7-fold shift in potency between the Wuhan-Hu-1 spike sequence and the B.1.1.7 spike variant (**Figure 6c**).

**Figure 6.**
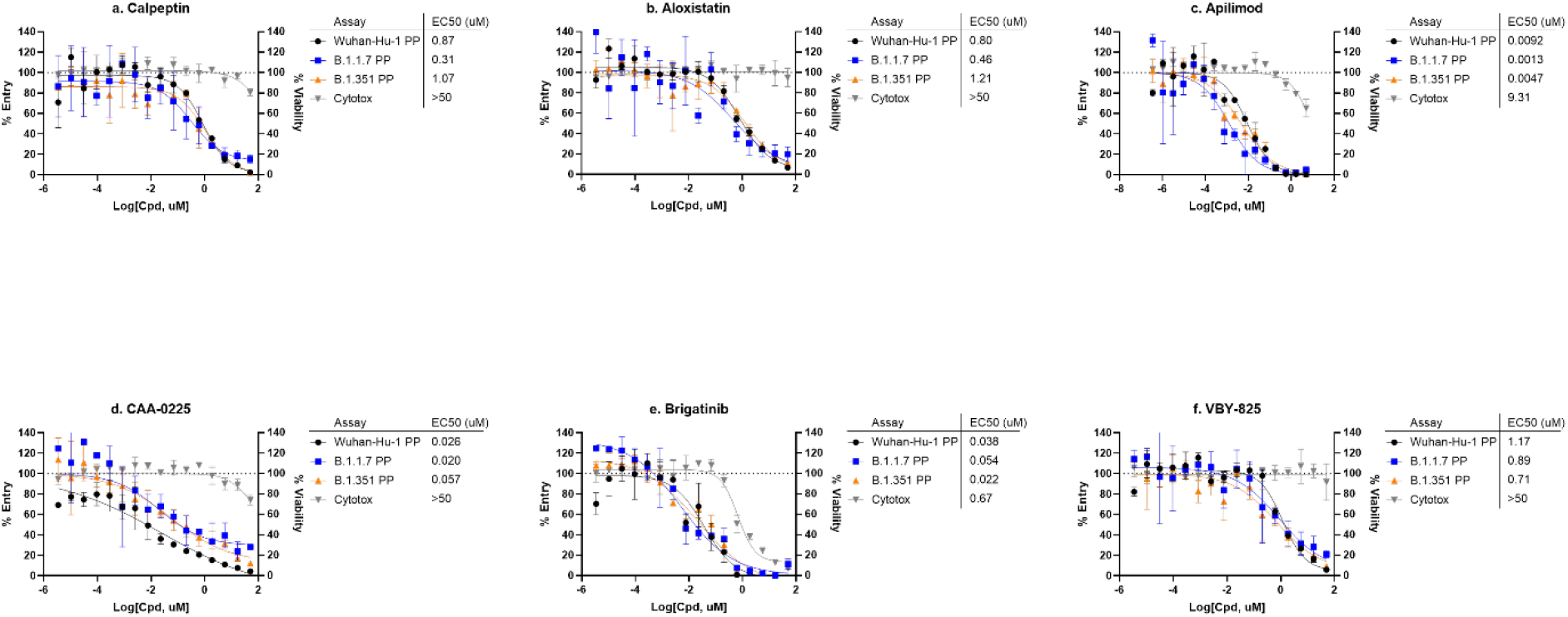
SARS-CoV-2-S variants PP assays. Concentration-response curves of (a) calpeptin, (b) aloxistatin, (c) apilimod, (d) CAA-0225, (e) brigatinib, and (f) VBY-825 in PP entry assays with Wuhan-Hu-1, B.1.1.7, and B.1.351 spike variants.

## Discussion

For highly pathogenic viruses such as Ebola, SARS, MERS, and SARS-CoV-2, BSL-2 assays such as pseudotyped viral particle assays and virus-like particle (VLP) assays are commonly used for HTS of compound collections for drug development ^4, 12^. The requirement of BSL-3 or BLS-4 facilities for live virus experiments could be a limiting factor for drug development. Therefore, here we report the design and development of an alternative BSL-2 containment level assay using SARS-CoV-2 PPs. The SARS-CoV-2 PPs consisted of SARS-CoV-2 spike protein incorporated in an enclosed bilayer membrane, packaged by MLV gag-pol core polyproteins, and enclosign luciferase reporter RNA. The incubation of SARS-CoV-2-S PPs with HEK293-ACE2 cells allows entry and expression of luciferase reporter gene in host cells that can be detected by luciferase reporter activity. Much like host cell entry of replicating SARS-CoV-2 virus, the entry of SARS-CoV-2-S PPs are mediated by spike protein interaction with host ACE2 receptor, cleavage of spike protein by host proteases to enable the triggering of conformational changes to the spike protein required for membrane fusion. These activities are dependent not only on interactions of spike protein with host cell receptor and proteases, but also on host cell endocytic machinery, and are therefore, highly cell type specific ^3^. Therefore, PP entry assays are phenotypic assays that can be used to screen multiple targets involved in viral cell entry.

This SARS-CoV-2-S PP entry assay is robust for compound screening in the 1536-well plate format, and is a good tool for HTS efforts for COVID-19 drug development. This assay can also be used to deconvolute mechanisms of action of anti-SARS-CoV-2 hits from live SARS-CoV-2 screens to identify compounds that act on viral entry versus viral replication. Viral pseudotyped assays have been commonly used to examine neutralization activity of antibodies and other biologics that block spike protein binding to the ACE2 receptor ^13^. However, unlike testing neutralizing antibodies, which have a clear target and mechanism of action, there is less certainty about the molecular target and the mechanism of inhibition when screening cell penetrant small molecules, which introduces additional mechanisms for false positives, such as inhibition of luciferase reporter expression. In our primary screen, some of the highly potent hits were known reverse transcriptase inhibitors (adefovir and tenofovir disoproxil fumarate) and integrase inhibitors (elvitegravir and dolutegravir), that were clearly functioning through inhibition of the MLV retroviral core proteins. These hits showed the same potency in the VSV-G PP entry counter assay, and were thus weeded out. However, use of VSV-G PP as a counter screen might be overly stringent, because while SARS-CoV-2-S and VSV-G have different host receptors and protease requirements, there are known entry mechanisms that are shared between the two glycoproteins. One such shared mechanism is binding to heparan sulfate proteoglycan to facilitate cell attachment ^14^. Future studies might feature viral-like particles (VLP), which directly tag viral packaging proteins with an enzyme protein reporter, and therefore eliminate the process of RNA reporter expression ^12^. However, successful SARS-CoV-2-S VLP generation with reporter enzyme has yet to be reported.

In the drug repurposing screen using this SARS-CoV-2-P PP entry assay, seven selective entry inhibitors were found including VBY-825, aloxistatin, apilimod, calpeptin, dichlorophen, brigatinib, and CAA-0225. Their activities against SARS-CoV-2 entry into host cells have been previously reported in the literature ^2, 15^. We also further examined their activity on the CPE of SARS-CoV-2 infection, resulting in confirmation of six compounds in the live virus assay, with the exception of dichlorophen (antiparasitic and antimicrobial agent) (Figure 4a, 5a). Four out of the six confirmed hits are known inhibitors of host proteases and block priming of spike protein after spike-ACE2 binding. VBY-825 is a potent cathepsin inhibitor originally developed for cancer therapy ^16^. Aloxistatin is a cysteine protease inhibitor developed for treatment of muscular dystrophy but the clinical trial was not successful ^17^. Calpeptin is a potent peptide inhibitor of cathepsin K and calpain ^18^. CAA-0225 is a potent peptide cathepsin L inhibitor ^19^. Brigatinib is a dual kinase inhibitor that targets mutant epidermal growth factor receptor (EGFR) and anaplastic lymphoma kinase (ALK) and is an approved drug for cancer therapy. Apilimod is an approved drug for autoimmune diseases, including rheumatoid arthritis and Crohn’s disease, through inhibitions of production of IL-12 and IL-23 and PIKfyve (a lipid kinase) ^20^. In addition, the antiviral activities of Apilimod against SARS-CoV-2, Lassa fever, and Ebola have been reported ^21, 22^. It is worth mentioning that cell entry of SARS-CoV-2 is highly cell type dependent ^3^. While our screen identified compounds that are more specific to the SARS-CoV-2-S mediated entry pathway over the VSV-G mediated pathway, any host cell directed inhibitors might show different potencies and/or efficacies in other cell types. For instance, the known cathepsin inhibitor hits (VBY-825, aloxistatin, calpeptin and CAA-0225), would not be active against SARS-CoV-2 entry in cells, which uses a TMPRSS2 protease dependent pathway instead of cathepsin dependent pathway, such as Calu-3 cells ^23^.

Similar to many RNA viruses, mutations in the RNA genome of SARS-CoV-2 occur frequently. Variants of concern (VOC) that either cause increased infectivity or vaccine escape due to multiple spike protein mutations are actively being tracked (https://www.cdc.gov/coronavirus/2019-ncov/variants/index.html). Several mutations have been reported in the SARS-CoV-2 spike protein that have raised some concern about the efficacy of current vaccines drugs ^24^. One advantage of the PP assay reported here is the ease of swapping spike protein plasmids, i.e. in SARS-CoV-2 PP production cells, the Wuhan-Hu-1 spike plasmid can be rapidly replaced with a plasmid containing any of the newly identified spike variant constructs. To illustrate this, we have generated PPs with mutations in spike protein from the B.1.1.7 and B.1.351 variants, and conducted pilot experiments to test the top entry inhibitor hits from the repurposing screen. Therefore, th SARS-CoV-2 PP entry assay reported here, with facile and rapid modifications, can be used to evaluate anti-SARS-CoV-2 efficacy of therapeutics (binding and entry inhibitors) on new variants with mutations in spike protein. It is a valid *in vitro* model for testing drugs against emergent SARS-CoV-2 variants with spike protein mutations.

## Data availability statement

The datasets presented in this study can be found in online repositories. The names of the repository/repositories and accession numbers can be found in Table 2 (PubChem AIDs 1645846, 1645845, 1645844, 1645847). Primary screen data can also be found at http://opendata.ncats.nih.gov. All other data are available upon request.

## Acknowledgements

This work was supported by the Intramural Research Program of National Center for Advancing Translational Sciences, Sciences, and National Institute of Child Health and Human Development, National Institutes of Health. This work was funded in part by the National Institutes of Health research grant R01AI35270.

## References

1. Shyr, Z. A.; Gorshkov, K.; Chen, C. Z.,; et al. Drug Discovery Strategies for SARS-CoV-2. J Pharmacol Exp Ther 2020, 375, 127–138.

2. Chen, C. Z.; Shinn, P.; Itkin, Z.,; et al. Drug Repurposing Screen for Compounds Inhibiting the Cytopathic Effect of SARS-CoV-2. Frontiers in Pharmacology 2021, 11.

3. Tang, T.; Bidon, M.; Jaimes, J. A.,; et al. Coronavirus membrane fusion mechanism offers a potential target for antiviral development. Antiviral Res 2020, 178, 104792.

4. Chen, C. Z.; Xu, M.; Pradhan, M.,; et al. Identifying SARS-CoV-2 entry inhibitors through drug repurposing screens of SARS-S and MERS-S pseudotyped particles. bioRxiv 2020.

5. Johnson, M. C.; Lyddon, T. D.; Suarez, R.,; et al. Optimized Pseudotyping Conditions for the SARS-COV-2 Spike Glycoprotein. Journal of Virology 2020, 94.

6. Millet, J. K.; Tang, T.; Nathan, L.,; et al. Production of Pseudotyped Particles to Study Highly Pathogenic Coronaviruses in a Biosafety Level 2 Setting. J Vis Exp 2019.

7. Millet, J. K.; Whittaker, G. R. Murine Leukemia Virus (MLV)-based Coronavirus Spike-pseudotyped Particle Production and Infection. Bio Protoc 2016, 6.

8. Xiao, T.; Lu, J.; Zhang, J.,; et al. A trimeric human angiotensin-converting enzyme 2 as an anti-SARS-CoV-2 agent. Nature Structural & Molecular Biology 2021, 28, 202–209.

9. Huang, R.; Zhu, H.; Shinn, P.,; et al. The NCATS Pharmaceutical Collection: a 10-year update. Drug Discov Today 2019, 24, 2341–2349.

10. Wang, Y.; Jadhav, A.; Southal, N.,; et al. A grid algorithm for high throughput fitting of doseresponse curve data. Curr Chem Genomics 2010, 4, 57–66.

11. Sun, X.; Roth, S. L.; Bialecki, M. A.,; et al. Internalization and fusion mechanism of vesicular stomatitis virus and related rhabdoviruses. Future Virology 2010, 5, 85–96.

12. Kouznetsova, J.; Sun, W.; Martinez-Romero, C.,; et al. Identification of 53 compounds that block Ebola virus-like particle entry via a repurposing screen of approved drugs. Emerg Microbes Infect 2014, 3, e84.

13. Nie, J.; Li, Q.; Wu, J.,; et al. Quantification of SARS-CoV-2 neutralizing antibody by a pseudotyped virus-based assay. Nature Protocols 2020, 15, 3699–3715.

14. Zhang, Q.; Chen, C. Z.; Swaroop, M.,; et al. Heparan sulfate assists SARS-CoV-2 in cell entry and can be targeted by approved drugs in vitro. Cell Discov 2020, 6, 80.

15. Riva, L.; Yuan, S.; Yin, X.,; et al. Discovery of SARS-CoV-2 antiviral drugs through large-scale compound repurposing. Nature 2020.

16. Elie, B. T.; Gocheva, V.; Shree, T.,; et al. Identification and pre-clinical testing of a reversible cathepsin protease inhibitor reveals anti-tumor efficacy in a pancreatic cancer model. Biochimie 2010, 92, 1618–1624.

17. Satoyoshi, E. Therapeutic Trials on Progressive Muscular Dystrophy. Internal Medicine 1992, 31, 841–846.

18. Tsujinaka, T.; Kajiwara, Y.; Kambayashi, J.,; et al. Synthesis of a new cell penetrating calpain inhibitor (calpeptin). Biochemical and Biophysical Research Communications 1988, 153, 1201–1208.

19. Takahashi, K.; Ueno, T.; Tanida, I.,; et al. Characterization of CAA0225, a Novel Inhibitor Specific for Cathepsin L, as a Probe for Autophagic Proteolysis. Biological & Pharmaceutical Bulletin 2009, 32, 475–479.

20. Ikonomov, O. C.; Sbrissa, D.; Shisheva, A. Small molecule PIKfyve inhibitors as cancer therapeutics: Translational promises and limitations. Toxicol Appl Pharmacol 2019, 383, 114771.

21. Baranov, M. V.; Bianchi, F.; van den Bogaart, G. The PIKfyve Inhibitor Apilimod: A Double-Edged Sword against COVID-19. Cells 2020, 10.

22. Hulseberg, C. E.; Feneant, L.; Szymanska-de Wijs, K. M.,; et al. Arbidol and Other Low-Molecular-Weight Drugs That Inhibit Lassa and Ebola Viruses. J Virol 2019, 93.

23. Hoffmann, M.; Mösbauer, K.; Hofmann-Winkler, H.,; et al. Chloroquine does not inhibit infection of human lung cells with SARS-CoV-2. Nature 2020, 585, 588–590.

24. Peacock, T. P.; Penrice-Randal, R.; Hiscox, J. A.,; et al. SARS-CoV-2 one year on: evidence for ongoing viral adaptation. Journal of General Virology 2021, 102.

